# A hybrid *in silico*/in-cell controller for microbial bioprocesses with process-model mismatch

**DOI:** 10.1101/2023.04.10.536196

**Authors:** Tomoki Ohkubo, Yuki Soma, Yuichi Sakumura, Taizo Hanai, Katsuyuki Kunida

**Affiliations:** Graduate School of Science and Technology, Nara Institute of Science and Technology, Ikoma, Nara, 8916-5, Japan; Laboratory for Synthetic Biology, Graduate School of Bioresource and Bioenvironmental Sciences, Kyushu University, W5-729, 744, Motooka, Nishi-ku, Fukuoka 819-0395, Japan; Data Science Center, Nara Institute of Science and Technology, Ikoma, Nara, 8916-5, Japan; School of Medicine, Fujita Health University, Toyoake, Aichi, 470-1192, Japan

**Keywords:** bioprocess, *Escherichia coli*, model-based optimization, structured model, synthetic biology

## Abstract

The optimization of bioprocess inputs using mathematical models is widely practiced. However, the mismatch between model prediction and the actual process [called process-model mismatch (PMM)] is problematic; when a large PMM exists, the process inputs optimized using the mathematical model in advance are no longer optimal for the actual process. In this study, we propose a hybrid control system that combines model-based optimization (*in silico* feedforward controller) and feedback controllers using synthetic genetic circuits integrated into cells (in-cell feedback controller) – which we named the hybrid *in silico*/in-cell controller (HISICC) – as a solution to this PMM issue. As a proof of concept for HISICC, we constructed a mathematical model of an engineered *Escherichia coli* strain for the isopropanol production process that was previously developed. This strain contains an in-cell feedback controller, and its combination with an *in silico* controller can be regarded as an example of HISICC. We demonstrated the robustness of HISICC against PMM by comparing the strain with another strain with no in-cell feedback controller in simulations assuming PMM of various magnitudes.

## Introduction

A primary goal of bioprocess engineering is to produce a larger amount of a desired product at higher rates from a smaller amount of raw materials (Villadsen, Nielsen, and Lidén 2011; Dochain 2008). To achieve this, it is necessary to control the behavior of microorganisms so that their capabilities can be harnessed to the fullest extent. Approaches to this can be broadly classified into two categories: computerized process control (*in silico* feedforward control) and autonomous feedback control by synthetic genetic circuits integrated in cells (in-cell feedback control) (Del Vecchio, Dy, and Qian 2016; Hsiao, Swaminathan, and Murray 2018; Khammash 2022).

*In silico* feedforward control is an approach widely used in industries where mathematical models are used to predetermine the optimal values of inputs to the process, such as temperature, pH, and substrate and inducer feeds (Penloglou et al. 2017; Majewski and Domach 1990; Novak et al. 2015; Khanna and Srivastava 2005; Horvat et al. 2013). *In silico* feedforward control comprises a sophisticated approach that maximizes product yield by predicting the future process state and managing the various tradeoffs that arise with respect to process inputs. However, one challenge is that the predetermined input values are no longer optimal in the actual process when there is a significant mismatch between the model and the actual process (process-model mismatch, PMM). One solution to the PMM problem is model predictive control (MPC), in which the state variables of the ongoing process are measured in real-time and fed back to the control inputs (Milias-Argeitis et al. 2016; Hafidi et al. 2008; Santos et al. 2012; Tebbani et al. 2010; Uhlendorf et al. 2012; Ashoori et al. 2009; Chang, Liu, and Henson 2016; Xiong and Zhang 2005; Mahadevan and Doyle 2003). Although MPC is widely used for process control, it can be unavailable if the model includes the intracellular concentrations of RNA, metabolites, or enzymes as state variables that are difficult to monitor online.

In-cell feedback control is a nascent approach that emerged from synthetic biology (Hartline et al. 2021). In the last two decades, various examples of synthetic genetic circuits have been reported, such as those designed to control cell density (Izard et al. 2015; Vignoni et al. 2013; You et al. 2004; Holtz and Keasling 2010), co-culture composition (Honjo et al. 2019; Gutiérrez Mena, Kumar, and Khammash 2022), or intracellular protein expression levels (Dunlop, Keasling, and Mukhopadhyay 2010; Harrison and Dunlop 2012; Gardner, Cantor, and Collins 2000; Hsiao et al. 2015; Zhang, Carothers, and Keasling 2012). Unlike *in silico* controllers, in-cell controllers can only provide simple feedback control, such as proportional control; sophisticated control to maximize future product yields is difficult for in-cell controllers. Conversely, it can detect intracellular RNAs, enzymes, and metabolites, which are difficult to monitor using process sensors or biochemical analyses and provide feedback on cell behavior *in situ*. Therefore, the *in silico* model-based controller and the in-cell feedback controller complement each other’s limitations.

To overcome the PMM problem of *in silico* controllers, we propose a hybrid control strategy that combines a high-level *in silico* feedforward controller and a low-level in-cell controller (hybrid *in silico*/in-cell controller, HISICC) (**Fig. 1**). When the actual process state deviates from the prediction by the *in silico* controller due to PMM, the in-cell feedback controller senses it and corrects the cell behavior so that a decrease in the product yield is prevented. In this study, we demonstrate the concept of HISICC by applying the *in silico* model-based control to the isopropanol (IPA) production process using the two engineered *Escherichia coli* strains we reported on previously (Soma et al. 2014; Soma and Hanai 2015) as an example of the bioprocess. As described in detail in the Results, prediction error in cell growth is a critical PMM in this process, which leads to decrease in IPA yield. Since only one of these strains contains an in-cell feedback controller which detects cell density, this strain can be defined as an example of HISICC in combination with the *in silico* feedforward controller whereas the other strain cannot. We demonstrated the robustness of HISICC to PMM, namely prediction errors in cell growth, by comparing the two strains in terms of IPA yield in simulations where various magnitudes of PMM were assumed.

**Figure 1.**
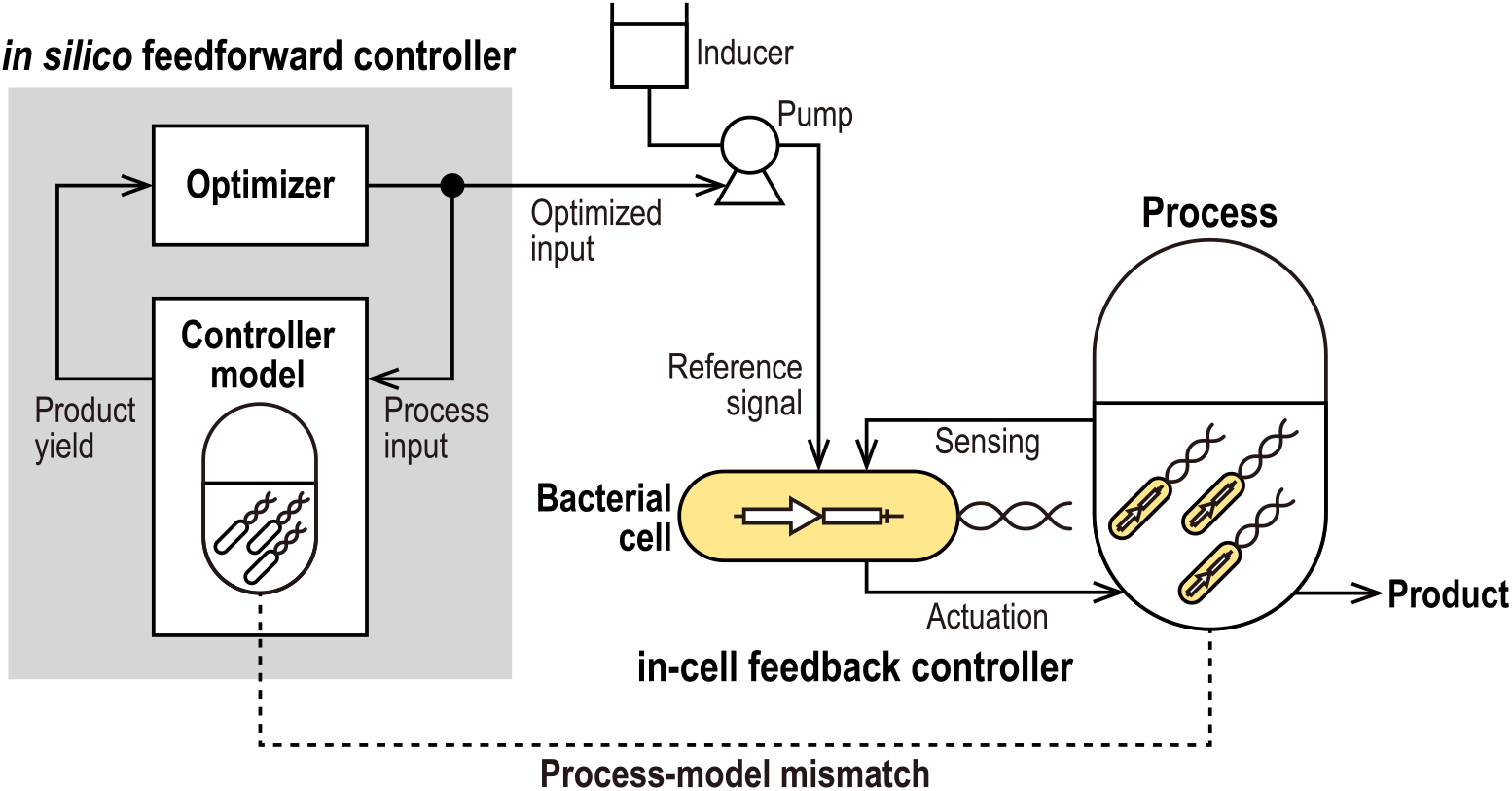
Conceptual diagram of the hybrid *in silico*/in-cell controller (HISICC). The *in silico* feedforward controller calculates the optimal control inputs for the process based on the controller model. An example of a control input is the inducer feed. The bacterial cells receive this control input as a reference signal and autonomously perform feedback control.

## Results

### IPA production process using two engineered strains

Prior to describing the details of mathematical modeling for the design of *in silico* feedforward controllers, we provide an overview of the IPA production process using the two engineered strains that we previously developed, TA1415 and TA2445 (Soma et al. 2014; Soma and Hanai 2015). In conventional IPA production processes, cell growth and IPA production compete for intracellular acetyl-CoA synthesized from the substrates. This competition needs to be balanced since an imbalance in the use of intracellular acetyl-CoA for either cell growth or IPA production results in reduced IPA yield. TA1415 has a genetic circuit called the metabolic toggle switch (MTS) that allows this competition to be managed by an external input of an inducer, isopropyl β-D-1-thiogalactopyranoside (IPTG). We designed an *in silico* feedforward controller that optimizes the IPTG input using a mathematical model of the strain. However, because TA1415 does not have an in-cell feedback controller, the combination of TA1415 and the *in silico* controller does not comprise an HISICC. In contrast, TA2445 has an in-cell feedback controller consisting of an MTS and another genetic circuit to detect cell density, termed quorum sensing. Owing to the in-cell feedback controller, TA2445 autonomously controls cell growth and IPA production in accordance with the external IPTG input as a reference signal. Therefore, the combination of TA2445 and an *in silico* feedforward controller designed for this strain can be considered an example of HISICC.

In both strains, the activated MTS stopped the synthesis of citrate synthase (the enzyme that mediates the reaction that initiates the TCA cycle) and simultaneously initiates the synthesis of a series of enzymes for IPA production, thereby achieving a changeover from cell growth to IPA production. The timing of MTS activation creates a tradeoff: if the MTS is activated too early, the IPA yield is low because the cells do not grow sufficiently; if the MTS is activated too late, the IPA yield is also low because extracellular nutrients are used up by cell growth, resulting in insufficient synthesis of a series of enzymes for IPA production.

In TA1415 cells, the MTS was activated by the addition of IPTG to the medium in the middle of the culture period (**Fig. 2A**). Therefore, the timing of IPTG addition can be defined as the input variable of the process to be optimized (**Fig. 2B**).

**Figure 2.**
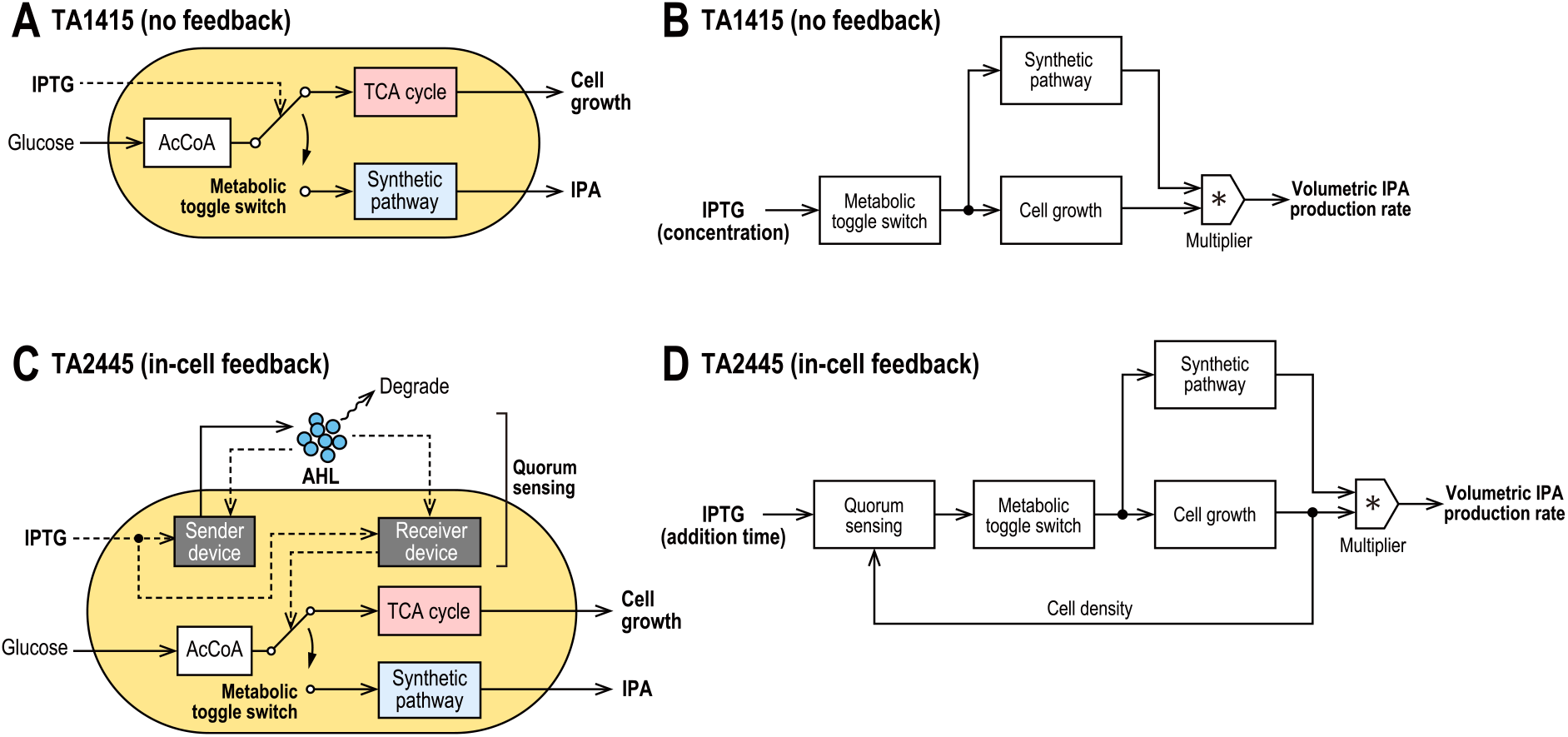
Genetic circuits of TA1415 and TA2445. (A) Genetic circuit of TA1415, in which the metabolic toggle switch (MTS) changes the flow of intracellular Acetyl-CoA (AcCoA) from the TCA cycle to the synthetic pathway for isopropanol (IPA) production; the MTS is activated when IPTG is added to the medium. (B) Block diagram showing the control structure of TA1415, in which the MTS changes the expression level of the synthetic pathway and cell growth. The volumetric production rate of IPA is proportional to the product of cell density and the expression level of the synthetic pathway. (C) Genetic circuit of TA2445 with the sender device to secrete and the receiver device to detect AHL, which realize quorum sensing collectively; the MTS is activated when the receiver device detects an increased extracellular concentration of AHL due to cell growth. (D) Block diagram showing the control structure of TA2445. Quorum sensing provides feedback of increased cell density to the MTS, the sensitivity of which depends on IPTG concentration in the medium.

TA2445 has an additional genetic circuit for quorum sensing that detects cell density to activate the MTS, as described in the Introduction (**Fig. 2C**). The circuit is composed of an intercellular messenger called an acylated homoserine lactone (AHL) and genetic devices that send or receive it. As the cell density increases, so does the AHL concentration in the medium. When the AHL concentration reaches a certain level, the receiver device detects AHL and activates the MTS. The sender and receiver devices utilize the same promoter, which responds to both IPTG and AHL. This allows the sensitivity of quorum sensing to be tuned by varying the extracellular concentration of IPTG. Thus, IPTG concentration can be defined as the input variable of the process to be optimized (**Fig. 2D**); if IPTG concentration is too high, quorum sensing becomes too sensitive, and the MTS is activated too early. Conversely, if the IPTG concentration is too low, quorum sensing becomes too insensitive, and the MTS is activated too late or is not activated at all.

### Mathematical modeling

#### TA1415 model

The TA1415 model is based on a two-compartment model, which is a type of structured model constructed by Williams that divides the cells into two compartments: XA and XG (Williams 1967). The XA compartment represents the active part of cells directly involved in cell growth, including RNA, ribosomes, and small metabolites such as amino acids. On the other hand, the XG compartment represents an inactive part that is not directly involved in cell growth, including DNA, proteins, and cell membranes. XA is produced from the extracellular substrate S, and XG is produced from XA. Since the amount of XG per cell is nearly constant, it can be considered proportional to the cell density. Williams’ two-compartment model, although quite simple, can explain the lag phase as well as the experimental fact that cell growth continues for a period after removal of the substrate from the medium in the middle of the log phase. In the simulation, no additional XA is produced after substrate removal, while XG is produced until the XA present in the cells is exhausted.

When IPTG is fed to TA1415 cells, cell growth slows but does not immediately stop (Soma et al. 2014). This behavior is similar to that observed after substrate removal during the log phase, as described above. This suggests that when the MTS stops the TCA cycle, cells store materials for cell growth, such as amino acids, which can be used to continue cell growth. Thus, to model the MTS, we extended Williams’ two-compartment model to a three-compartment model with an additional compartment, E, representing a series of enzymes for IPA production (**Fig. 3A, Tables 1 and 3**).

**Figure 3.**
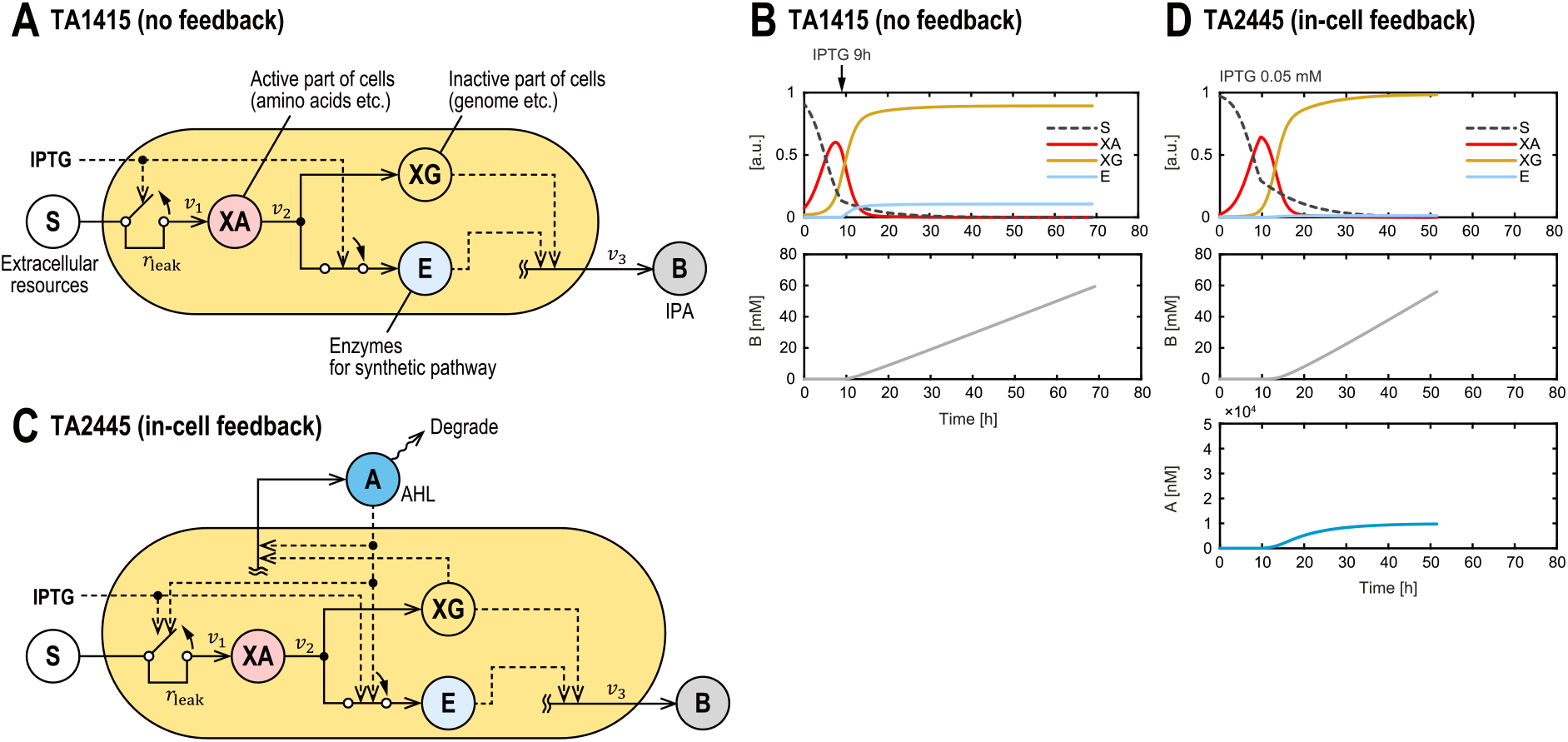
The three-compartment models of the two strains. (A) TA1415 model. Active compartment XA is synthesized by the TCA cycle from S, which represents extracellular resources; from XA, inactive compartment XG and E, a series of enzymes on the synthetic pathway for IPA production, are synthesized. When activated, the MTS stops synthesizing XA and initiates synthesis of E. The production rate of IPA, represented by B is proportional to XG and E. (B) Dynamics of state variables for TA1415 model, with IPTG added at 9 h to activate the MTS. (C) TA2445 model. A, which represents extracellular AHL, increases with cell growth. The MTS is activated when A reaches a certain level that depends on extracellular IPTG concentration. (D) Dynamics of state variables for TA2445 model at IPTG concentration of 0.05 mM.

**Table 1.** Equations for the three-compartment model of TA1415

| Description | Formula |  |
| --- | --- | --- |
| Reaction rates | $v_1 = k_1 S(X_A + X_G + E)(1 - (1 - r_{\text{leak}})u)$ | 1 |
| | $v_2 = k_2 X_A(X_G + E)$ | 2 |
| | $v_3 = k_3 \frac{E}{E + K_E} X_G$ | 3 |
| State equations | $\frac{dS}{dt} = -v_1$ | 4 |
| | $\frac{dX_A}{dt} = v_1 - v_2$ | 5 |
| | $\frac{dX_G}{dt} = v_2(1 - a_E u)$ | 6 |
| | $\frac{dE}{dt} = v_2 a_E u$ | 7 |
| | $\frac{dB}{dt} = v_3$ | 8 |
| Observation equations | $y_1 = N_m X_G$ for cell density | 9 |
| | $y_2 = B$ for IPA concentration | 10 |

In our three-compartment model, XA is produced from S by the TCA cycle (Reaction 1), and XG and E are synthesized from XA (Reaction 2). The production rate of B, which represents IPA, was proportional to both XG and E (Reaction 3). We introduced a saturation constant, *K*_*E*_ so that the rate of Reaction 3 was saturated with respect to E. When MTS is not activated, Reaction 1 proceeds, and only XG is produced in Reaction 2. When MTS is activated, Reaction 1 stops, and E is produced in addition to XG in Reaction 2 (**Fig. 3B**). However, because of some leakage in the promoter, Reaction 1 does not stop completely, and a small amount of XA continues to be produced thereafter.

#### TA2445 model

The TA2445 model is based on the three-compartment model of TA1415, with the addition of a part of the quorum sensing model constructed by You et al., in which AHL secreted from cells is degraded in a first-order reaction (**Fig. 3C, Tables 2 and 3**) (You et al. 2004). Simulated trajectories of each compartment at IPTG concentration of 0.05 mM are shown in **Fig. 3D**. The responses of the sender and receiver devices to IPTG and AHL were modeled using data from another *E. coli* strain we previously developed, TA2393 (Soma and Hanai 2015). TA2393 has a GFP gene on a plasmid downstream of the same promoter as the sender and receiver devices of TA2445. By using TA2393, we measured the promoter response to IPTG and AHL in terms of fluorescence intensity. The response curve of the promoter was fitted to the Hill equation to obtain the values of its four parameters: the dissociation constants and Hill coefficients for IPTG or AHL (**Fig. 4**, *Equation 14*). Approximating the promoter response using the product of the Hill equations for AHL and IPTG was reported in a previous study using similar promoters (Sekine et al. 2011).

**Table 2.** Equations for the three-compartment model of TA2445

| Description | Formula |  |
| --- | --- | --- |
| Reaction rates | $v_1 = k_1 S(X_A + X_G + E)(1 - (1 - r_{\text{leak}})z)$ | 11 |
| | $v_2 = k_2 X_A(X_G + E)$ | 12 |
| | $v_3 = k_3 \frac{E}{E + K_E} X_G$ | 13 |
| Promotor response to IPTG and AHL | $z = \frac{A^{n_A}}{A^{n_A} + K_A^{n_A}} \frac{u^{n_u}}{u^{n_u} + K_u^{n_u}}$ | 14 |
| State equations | $\frac{dS}{dt} = -v_1$ | 15 |
| | $\frac{dX_A}{dt} = v_1 - v_2$ | 16 |
| | $\frac{dX_G}{dt} = v_2(1 - a_E z)$ | 17 |
| | $\frac{dE}{dt} = v_2 a_E z$ | 18 |
| | $\frac{dB}{dt} = v_3$ | 19 |
| | $\frac{dA}{dt} = k_A X_G z - d_A A$ | 20 |
| Observation equations | $y_1 = N_m X_G$ for cell density | 21 |
| | $y_2 = B$ for IPA concentration | 22 |

**Table 3.** State and input variables for the two models

| Symbol | Initial value for<br>TA1415 | Initial value for<br>TA2445 | Unit | Description |
| --- | --- | --- | --- | --- |
| **for TA1415 and TA2445** |  |  |  |  |
| $S$ | $1 - X_A(0) - X_G(0)$ | $1 - X_A(0) - X_G(0)$ | dimensionless | Extracellular resources |
| $X_A$ | estimated | estimated | dimensionless | Active compartment |
| $X_G$ | 0.02 | 0.005 | dimensionless | Inactive compartment |
| $B$ | 0 | 0 | mM | IPA concentration |
| $E$ | 0 | 0 | dimensionless | Enzymes for IPA production |
| **for TA1415** |  |  |  |  |
| $u$ | — | — | dimensionless | Normalized IPTG concentration |
| **for TA2445** |  |  |  |  |
| $A$ | — | 0.01 | nM | AHL concentration |
| $u$ | — | — | mM | IPTG concentration |

**Table 4.** Parameters for the two models estimated with all datasets

| Symbol | Value for<br>TA1415 | Value for<br>TA2445 | Unit | Description |
| --- | --- | --- | --- | --- |
| **for TA1415 and TA2445** |  |  |  |  |
| $k_1$ | 0.4772 | 0.4526 | /h | Production rate parameter for Reaction 1 |
| $k_2$ | 0.9089 | 1 | /h | Production rate parameter for Reaction 2 |
| $k_3$ | 1.2418 | 2 | mM/h | Production rate parameter for Reaction 3 |
| $r_{leak}$ | 0.2777 | 0.0607 | dimensionless | Leak ratio of Reaction 1 |
| $a_E$ | 0.1508 | 0.0190 | dimensionless | Ratio of XA allocated to the synthesis of E |
| $K_E$ | 0.0081 | 0.0034 | dimensionless | Saturation coefficient for E in production rate of B |
| $X_A(0)$ | 0.0572 | 0.0240 | dimensionless | Initial value of XA. Ten times of $X_G(0)$ was set as the upper limit of estimation. |
| $N_m$ | 5.4651 | 5.6799 | OD600 | Carrying capacity |
| **for TA2445** |  |  |  |  |
| $k_A$ | — | 1.9137E+3 | nM/h | AHL production rate parameter |
| $d_A$ | — | 0.1489 | /h | AHL degradation rate parameter |
| $K_A$ | — | 1.596 | nM | Dissociation constant for AHL<br>(estimated with TA2393 data) |
| $K_u$ | — | 0.02268 | mM | Dissociation constant for IPTG<br>(estimated with TA2393 data) |
| $n_A$ | — | 1.752 | dimensionless | Hill coefficient for AHL<br>(estimated with TA2393 data) |
| $n_u$ | — | 1.597 | dimensionless | Hill coefficient for IPTG<br>(estimated with TA2393 data) |

**Figure 4.**
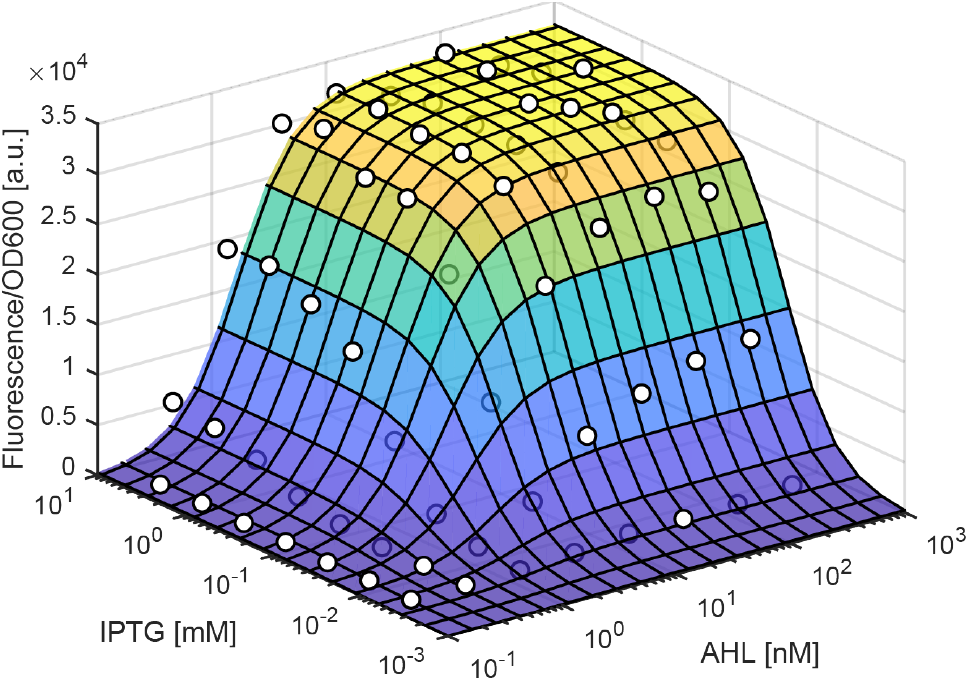
Response to IPTG and AHL of the promoters used in the sender and receiver devices of TA2445. Experimental data (white dots) of the *Escherichia coli* strain TA2946, which has a plasmid with the GFP gene downstream of the same promoter, was fitted using Hill equation.

### Model simulation and validation

The TA1415 and TA2445 models were trained using experimental data on the IPA production process obtained in previous studies (Soma et al. 2014; Soma and Hanai 2015). The details of the experimental data are described in the Materials and methods section. Both trained models fit the experimental data closely (**Fig. 5**). This indicates that, despite their simple structure, our models capture the dynamics of cell growth and IPA production of the two strains in response to various IPTG inputs. Additionally, we used the hold-out validation method to ensure that the two trained models did not overfit the training data (**Fig. 6**). The details of the validation method are described in the Materials and methods section. The coefficients of determination *R*^2^ were above 0.5 for all test data, indicating that both models have adequate generalization performance within the range of IPTG input values of the training data. The slightly lower *R*^2^ values for IPA concentration than for cell density (OD600) may be because the three-compartment model does not represent the slowdown of IPA production rate due to substrate depletion.

**Figure 5.**
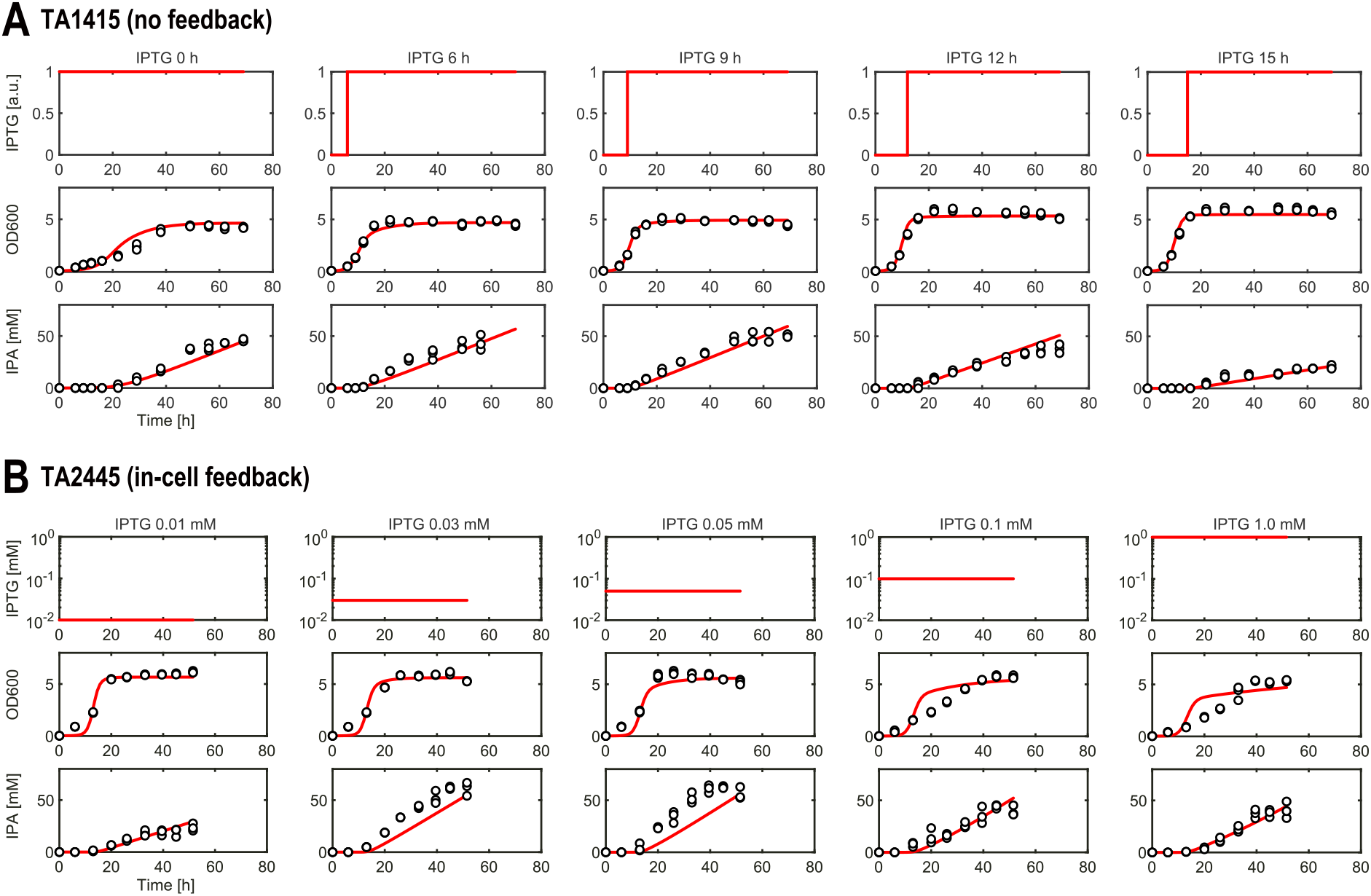
Simulations of the IPA production process for various IPTG inputs using the models of the two strains. Both models were trained using all datasets. White dots represent experimental data and red lines represent simulation results. (A) for TA1415. (B) for TA2445.

**Figure 6.**
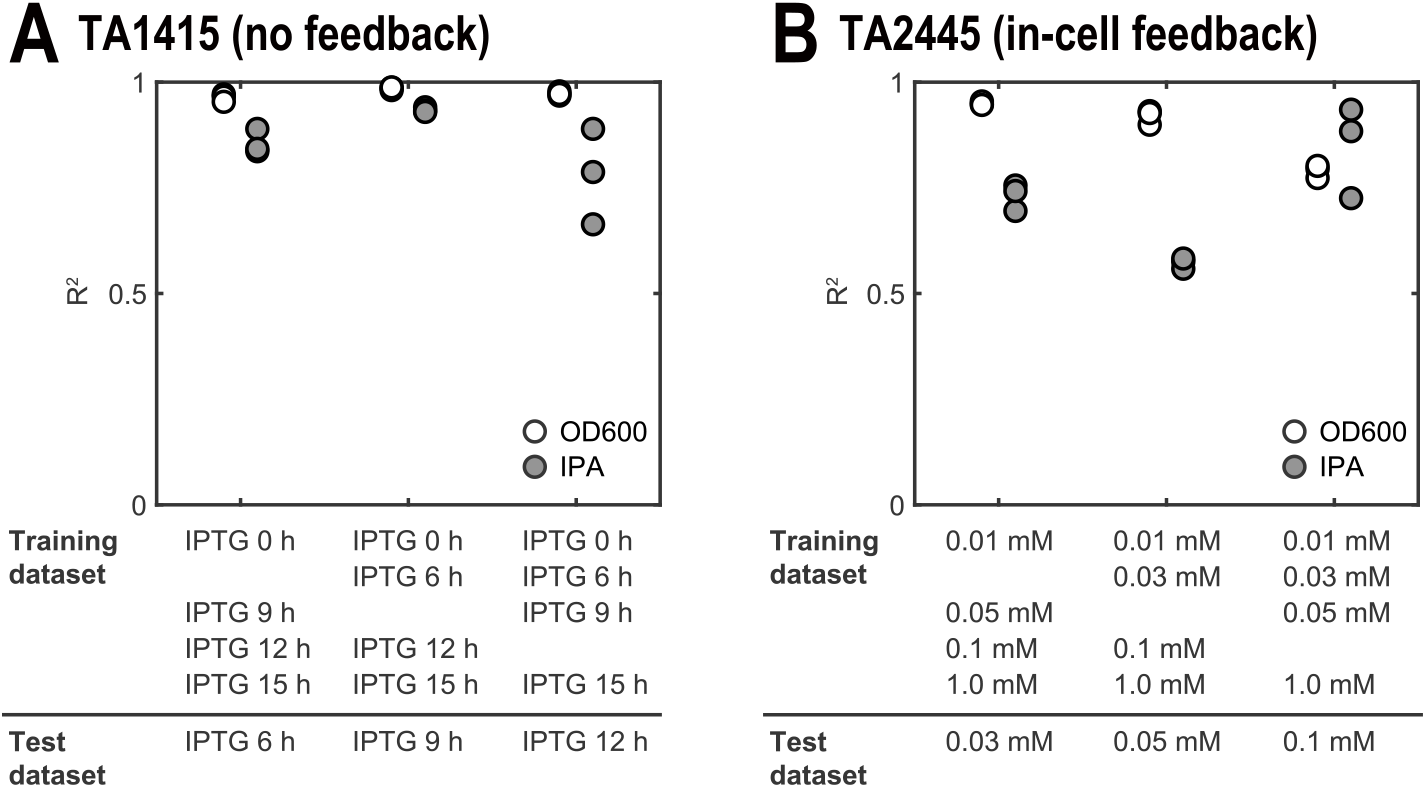
Hold-out validation of the models of the two strains, using a dataset subjected to one of five IPTG input conditions as a test dataset. (A) for TA1415. (B) for TA2445.

### Model-based input optimization

To demonstrate the optimal control by the *in silico* feedforward controller, we optimized the IPTG input variables to maximize the IPA concentration at the end of the culture using the two models. The timing of IPTG addition for TA1415 (**Fig. 7A**) and the concentration of IPTG for TA2445 (**Fig. 7B**) were optimized. The feasible region for IPTG input was defined as 0–15 h for TA1415 and 0.01–1.0 mM for TA2445. In addition, to visualize the overall distribution of IPA yield over the range of feasible IPTG input values, we comprehensively simulated the models within this range. The model predictions captured the experimental trends, which had a single peak, indicating that our models successfully reproduced the tradeoff in the IPA production process with both strains.

**Figure 7.**
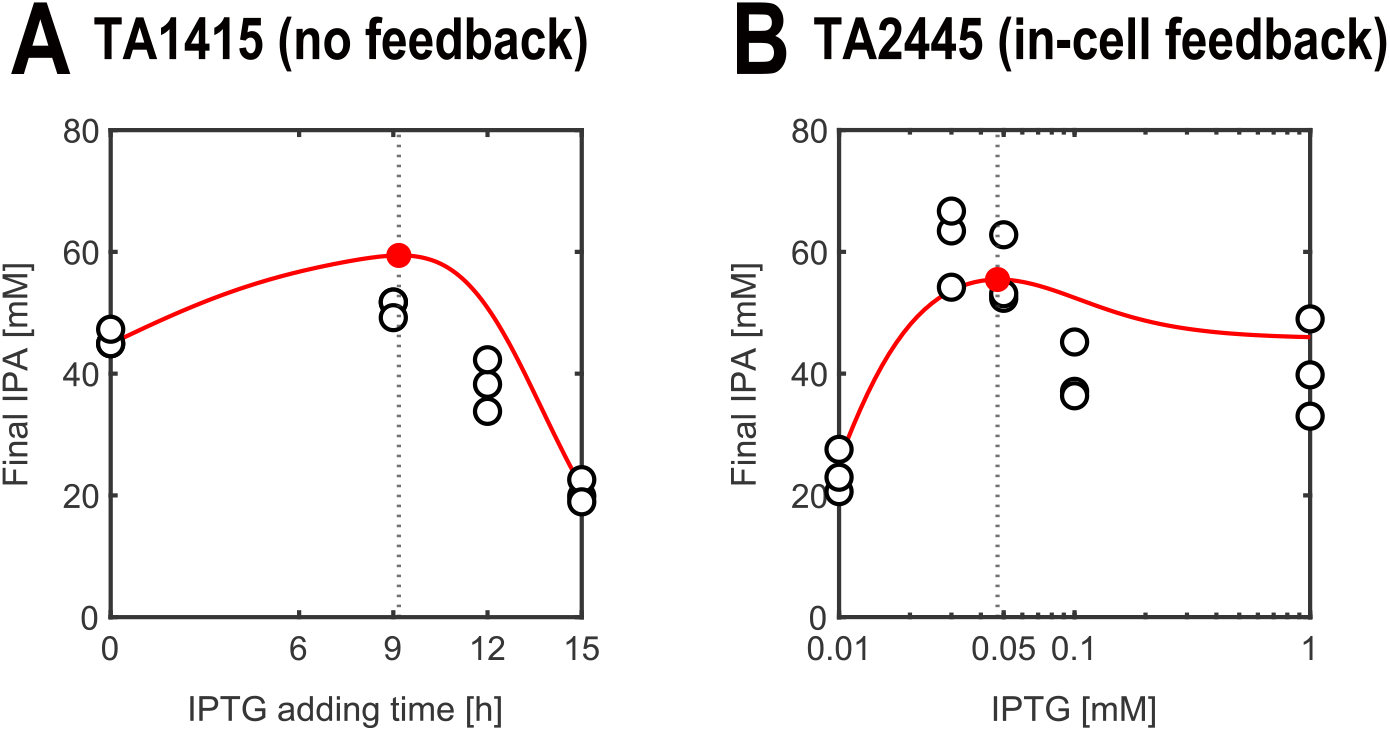
Model-based optimization of IPTG inputs to maximize IPA yield. White and red dots represent experimental data and optimized values, respectively, and red lines represent results of the exhaustive simulations. (A) for TA1415. (B) for TA2445.

### Controller performance against PMM

To evaluate the robustness of HISICC against PMM, we compared IPA yields of the two strains in the simulation. The simulation assumed that the growth rate of both strains was faster or slower than that predicted by the *in silico* feedforward controller. For two parameters (*k*_1_ and *k*_2_) which determine the cell growth rate, we set different values for the actual process compared with the controller model. Setting 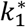 and 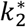 as the values for the actual process. In other words, when 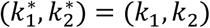 there is no PMM. The IPA yield maximized in the absence of PMM is referred to as the nominal yield. We performed exhaustive simulations using different combinations of 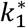 and 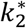 values to obtain the IPA yield (**Fig. 8**). First, for both strains, when the growth rate was slower than prediction (namely 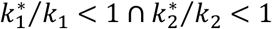), IPA yields were lower than the nominal yields. This was apparently because cell density did not increase sufficiently during the fixed culture period. However, when the cell growth was faster than that predicted by the controller model, the two strains resulted in different IPA yields. In the case of TA1415 (which did not contain an in-cell feedback controller), the addition of IPTG was delayed relative to the truly optimal timing, resulting in a lower IPA yield than the nominal yield (**Fig. 8A**). By contrast, in the case of TA2445, which contains an in-cell feedback controller, cells can autonomously adjust the timing of MTS activation earlier, resulting in suppression of the decrease in IPA yield (**Fig. 8B**). These results indicate that within HISICC, the in-cell feedback controller can support the *in silico* feedforward controller to prevent it from being disturbed by the PMM.

**Figure 8.**
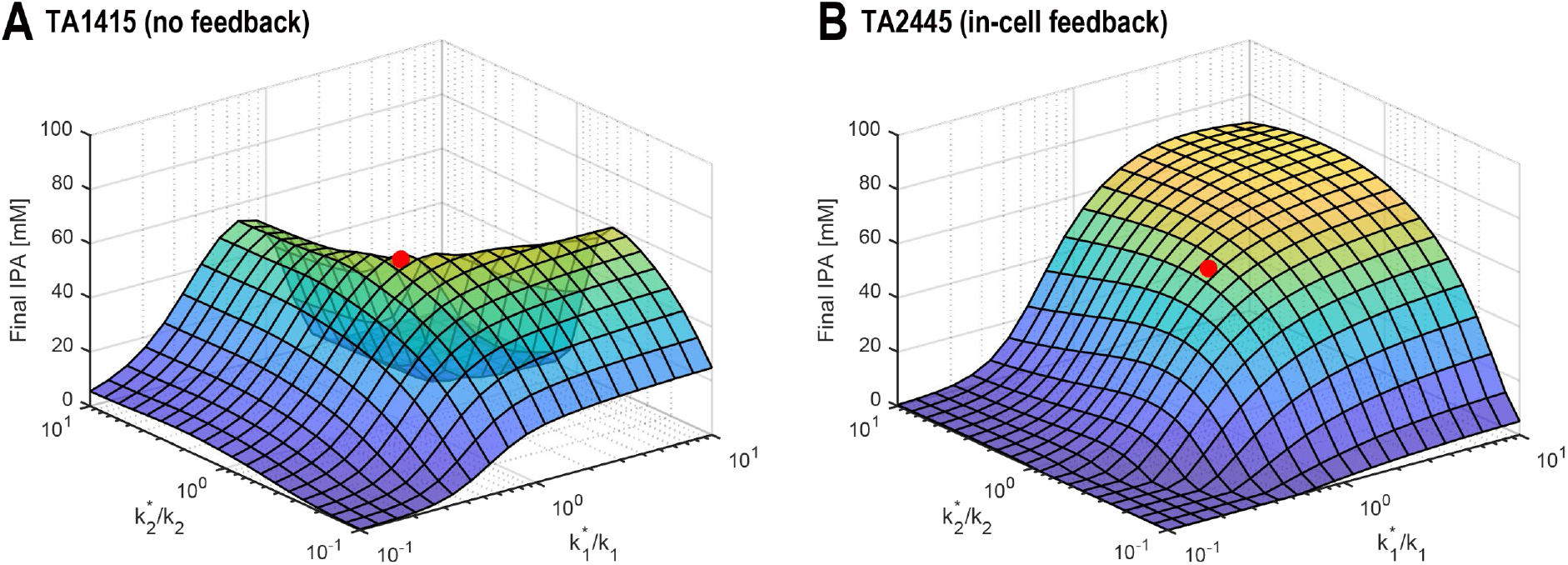
Variation of IPA yields with respect to PMM. The x-axis and y-axis represent the ratio of parameter values in the controller model and ones in the actual process for the two parameters that determine the cell growth rate. Red dots represent the nominal IPA yield (i.e., IPA yields maximized when the cell growth rate was defined equal between the controller model and in the actual process). (A) for TA1415. (B) for TA2445.

## Discussion

In this study, we proposed HISICC, a hybrid control system in which a high-level model-based controller provides a reference signal to a low-level in-cell feedback controller by means of the inducer concentration to suppress the performance deterioration caused by the PMM. We then performed a proof-of-concept of HISICC in the IPA production process with two *E. coli* strains that contain the MTS. Only one of these strains can be combined with the *in silico* feedforward controller to form a HISICC because it has an in-cell feedback controller that detects cell density (quorum sensing) to activate the MTS. We hypothesized that owing to the in-cell feedback controller, this HISICC can correct the timing of the MTS activation based on quorum sensing to prevent IPA yield decrease due to PMM of cell growth rate. To prove the hypothesis, first, mathematical models of the two strains were constructed to design an *in silico* feedforward controller. The constructed models are based on a previously reported two-compartment model. We used the experimental data from the IPA production culture to estimate the values of the parameters included in these models. Although the constructed models had simple structures, they captured the dynamics of cell growth and IPA production in response to various IPTG inputs. Both models showed excellent prediction performances for the experimental data in the hold-out validation. The validated models were then used to evaluate the robustness of HISICC against PMM. Finally, we compared the IPA yield between the two strains using simulations in which the model predictions and actual cell growth rates were assumed to be different. The results showed that, as we hypothesized, when cell growth is faster than expected by the *in silico* controller, the strain equipped with the in-cell feedback controller can prevent a decrease in IPA production. On the other hand, the strain without the in-cell controller cannot prevent a decrease in IPA production, which demonstrates the effectiveness of HISICC.

The cell density of the strain used in this proof-of-concept study was measured to autonomously activate the MTS. Since the cell density can be easily measured using a standard spectrophotometer instead of an in-cell feedback controller, it is easy to suppress the influence of the PMM by combining a low-level feedback controller using a spectrophotometer with a high-level model-based controller. However, as noted in the Introduction, in many microbial processes, the optimization of process inputs involves intracellular concentrations of mRNA, proteins, metabolites, or products. In such cases, few biochemical analysis methods are applicable for feedback control of the process because of their long turnaround times. Bacteria-based processes require particularly short turnaround times due to rapid cell growth. We believe that the HISICC proposed in this study can be a solution to the PMM problem when the ongoing monitoring of the process state is challenging using conventional hard sensors or biochemical analysis methods.

Furthermore, we must note a few points regarding the three-compartment models that we constructed. First, the state variables included in these models are approximate and difficult to interpret as concentrations of specific substances. Williams discussed the same issue in his two-compartment model (Williams 1967). In particular, substrate S in our models does not correspond explicitly to the glucose concentration in the medium, but rather abstractly represents the total extracellular resources consumed for cell growth and enzyme synthesis, including nutrients such as sugars and nitrogen sources, accumulation of waste products, and pH shifts. These abstractions allow our models to capture the dynamics of cell growth and IPA production in response to various IPTG inputs, while maintaining very simple structures. Secondly, as mentioned in the model simulation and validation section, our model does not account for the slowdown in the production rate of IPA due to substrate depletion at the end stage of culture, as in the model reported by Dunlop *et al*. (Dunlop, Keasling, and Mukhopadhyay 2010). This approximation would have resulted in a higher yield of IPA than the nominal yield in the simulation which assumed that the actual cell growth was faster than that predicted by the *in silico* controller with TA2445 (**Fig. 8B**). Thus, the increase in the IPA yield owing to faster cell growth was negligible. However, we believe that this approximation does not affect our argument that HISICC prevents the reduction in IPA yield due to PMM.

## Materials and methods

### Experimental data

The experimental data from the IPA production cultures used in this study to train and validate the models for the two engineered *E. coli* strains, TA1415 and TA2445, were obtained from two previously published studies (Soma et al. 2014; Soma and Hanai 2015). Here, we provide a brief description of the IPA production culture experiments. For both strains, seed cultures were grown overnight in 3 mL of M9 minimal medium supplemented with 10 g/L glucose, 1 g/L casamino acids, and 10 ppm thiamine hydrochloride at 37°C on a rotary shaker at 250 rpm. IPA production cultures were initiated with 1% (v/v) inoculation from the seed culture and grown in 20 mL of M9 minimal medium supplemented with 20 g/L glucose, 1 g/L casamino acids, and 10 ppm thiamin hydrochloride at 30°C on a rotary shaker at 250 rpm. Cell density (OD600) and IPA concentration were measured routinely during culture. For TA1415, the culture duration was 69 h. In the middle of the culture, 0.1 mM IPTG (concentrated enough to activate the MTS) was added at five different timepoints (0, 6, 9, 12, and 15 h). Three flasks were cultured for each addition of IPTG. For the TA2445 cells, the culture duration was 51.5 h. At the beginning of the culture period, different concentrations of IPTG (0.01, 0.03, 0.05, 0.1, or 1.0 mM) were added to the medium to tune the in-cell feedback controller. Three flasks were cultured for each addition of IPTG.

### Parameter estimation

MATLAB/Simulink 2022a was used for model construction and simulation. In the modeling of TA2445, Curve Fitting Toolbox was used to approximate the promoter response to AHL and IPTG using the Hill equation, as described in the mathematical modeling and simulation section. Simulink Design Optimization was used to estimate the other model parameters. The parameter values were chosen to minimize the sum of the squared errors between the model predictions and the measured data, as shown in *Equations 23* and *24*. Errors were normalized to the maximum values of measurements in the same culture. In these equations, *V* represents the objective function for optimization. Vector *θ* and 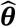 represent the model parameters and estimated values for them, respectively. *y* and *ŷ* represent the measured and predicted process outputs, respectively. ũ represents the IPTG input (addition time for TA1415 and concentration for TA2445). The subscripts *i, j*, and *k* represent the process output index (*i* = 1 for cell density and *i* = 2 for IPA concentration), culture flask index, and measurement time index, respectively.

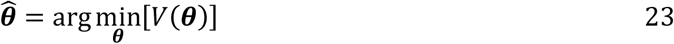

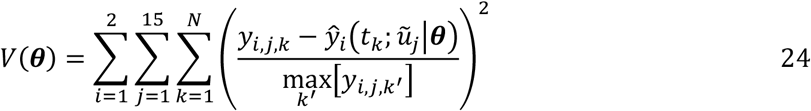

The *lsqnonlin* command was used for optimization. The trust region method was selected as the optimization algorithm for the command. A scaling factor was specified for each parameter to prevent those with large absolute values from excessively influencing the overall parameter estimation.

### Model validation

We validated that the constructed models correctly predicted the cell density and IPA concentration in response to different IPTG input values using the hold-out method. For each round of validation, one IPTG input value was selected from the five experimental values, excluding the maximum and minimum values. The experimental dataset from one of the three flasks to which the selected input condition was applied was defined as the test dataset. The datasets from the remaining 12 flasks were collectively defined as the training datasets. For each round of validation, the coefficient of determination *R*^2^ was calculated for the cell density or IPA concentration as follows:

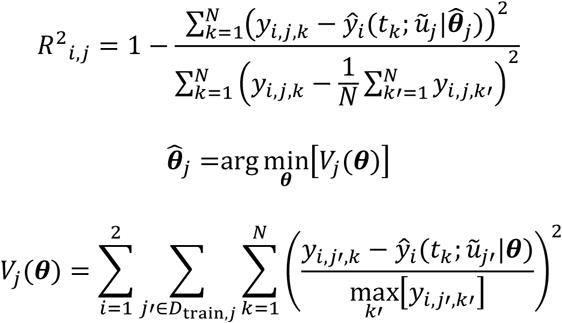

The subscripts *j* = 4,5, ⋯, 12 represents the flasks selected for the test dataset. The subscript set *D*_train,*j*_ represents the set of flasks selected for the training dataset.

### Model-based input optimization

Simulink Design Optimization was used to optimize the IPTG input. The optimal value *ũ*_*opt*_ was chosen to maximize the IPA concentration at the end of the culture, as shown in *Equations 25* and *26*. The culture duration *t*_*N*_ in the simulation was defined as 69 h and 51.5 h for TA1415 and TA2445, respectively, as in the experiments.

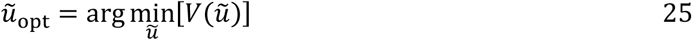

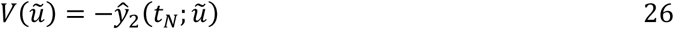

## Data availability

The datasets and computer code used in this study are available at GitHub (https://github.com/kkunida/202304_Ohkubo_bioRxiv.git).

## Acknowledgements

We thank Editage (www.editage.com) for English language editing. This study was supported by the Next Generation Interdisciplinary Research Project of Nara Institute of Science and Technology (NAIST).

